## Supplementary figures and images for "The Hsf1-sHsp cascade has pan-antiviral activity in mosquito cells"

### Fig S1

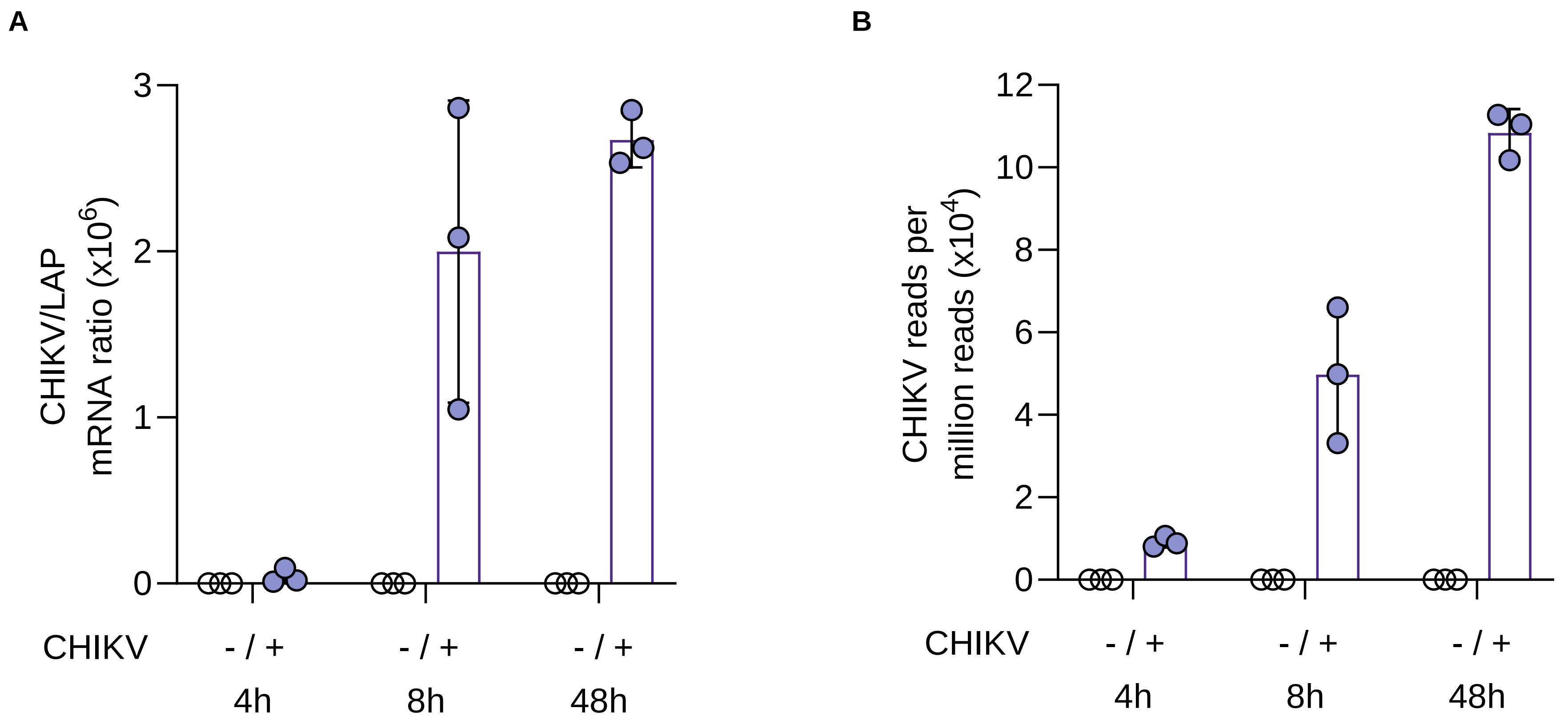
